# Decomposition of L-glutamine and accumulation of ammonium in cell culture media inhibit infectivity of influenza viruses

**DOI:** 10.1101/2024.06.21.600004

**Authors:** Nathanael B. Kegel, Andreas Kaufmann, Mikhail Matrosovich, Stefan Bauer, Jens Dorna

## Abstract

Cationic lysosomotropic molecules such as ammonium salts and chloroquine inhibit influenza A virus (IAV) infection in cell culture by counteracting endosomal acidification and hampering viral-endosomal fusion. Here, we studied the effects of storage of L-glutamine-supplemented cell culture media on accumulation of ammonium and inhibition of IAV infection. The storage-related inhibitory effect was observed in the case of DMEM and OptiMEM media, but not RPMI medium, and was more pronounced for the IAV with pH-stable hemagglutinin. Our results highlight the importance to consider potential presence of virus-inhibiting lysosomotropic agents and/or their production in culture media with negative effect on influenza virus infection.

## 1. Introduction

Influenza A virus (IAV) is responsible for up to one billion human infections per year (Lambert and Fauci 2010). The virus belongs to the *Orthomyxoviridae* family and possesses a lipid envelope and eight segmented, single-stranded antisense RNA molecules. The combination of two major surface glycoproteins, hemagglutinin (HA) and neuraminidase (NA), determines the HA/NA antigenic subtypes, of which H1-H16 and N1-N9 are found in wild aquatic birds, the natural reservoir of the virus, and H1N1, H2N2 and H3N2 subtypes are associated with pandemic and seasonal infections in humans (Krammer et al. 2018).

The HA mediates IAV attachment to sialic acid-containing receptors on target cells. After endocytosis, decreasing pH in the endosomes triggers i) conformational transition of the HA and ii) dissociation of viral ribonucleoproteins (RNPs) from the matrix protein. These low-pH-dependent effects are required for HA-mediated fusion of viral and endosomal membranes and release of the RNPs into the cytosol (Gamblin et al. 2021; Carter and Iqbal 2024). The pH threshold of HA conformational transition determines the pH optimum of HA-mediated membrane fusion as well as resistance towards acidic and temperature-related environmental factors. This threshold typically falls within the range of pH 4.8 and 6.2, and partially depends on the virus host species, i.e. IAVs of wild ducks and humans often being more acid-stable compared to poultry-adapted IAVs (Russell et al. 2018).

Cationic lysosomotropic molecules such as ammonium salts, amantadine, and chloroquine, are long known to inhibit IAV infection in vitro (Jensen and Liu 1963; Yoshimura et al. 1982; Daniels et al. 1985). The inhibition depends, at least in part, on drug-mediated elevation of endosomal pH, which prevents HA conformational transition and viral-endosomal fusion. Based on this mechanism, the sensitivity of IAVs to inhibition by ammonium chloride was used as a correlate of conformational stability and optimal pH for fusion of the HA (Doms et al. 1986; Koerner et al. 2012; Gerlach et al. 2017).

Media used for cultivation of IAVs in mammalian cell culture typically include L-glutamine. This amino acid is a building block in protein biosynthesis, is a part of purine and pyrimidine metabolism, and an important energy source (Smith 1990). Metabolic conversion of L-glutamine by cells as well as its spontaneous decomposition in aqeuous solutions lead to generation of ammonium ions. The rate of the the reaction depends on many parameters, such as cell type and pH, composition, and temperature of the solution (Schneider 1996). Thus, Tritsch and Moore found that decomposition of L-glutamine in cell-free culture media follows first order kinetics with 10 % conversion within nine days at 4 °C and within one day at 37 °C (Tritsch and Moore 1962). As a result, culture media, which are typically supplemented with 2-4 mM of L-glutamine, can accumulate ammonium in equivalent concentrations (Heeneman et al. 1993), thus potentially reaching concentrations of ammonium sufficient to affect infectivity of IAVs. To address this possibility, we here studied how storage of three representative L-glutamine-containing cell culture media affects the infectivity of two IAV strains differing by the conformational stability of their HA. Our results highlight the importance to consider potential presence of virus-inhibiting lysosomotropic agents in cell culture media used for growing and analyses of influenza viruses in vitro.

## Material & Methods

### 2.1 Cells

MDCK cells were propagated in DMEM (Pan BioTech, P04-036001) supplemented with 2 mM L-glutamine (Gibco), 100 IU/ml penicillin-streptomycin (Gibco) and 10% foetal calve serum (Gibco). The cells were passaged twice per week in T75 plastic flasks and maintained at 37 °C,7.5 % CO_2_.

### 2.2 Viruses

The 2:6-recombinant IAVs rHK-wt and rHK-17R, generated using eight-plasmid reverse genetics, were described previously (Gerlach et al. 2017). The viruses shared six gene segments of the laboratory strain A/Puerto Rico/8/1934 (PR8) and HA/NA segments of A/Hong Kong/1/1968 (H3N2). rHK-wt contained the original HA of A/Hong Kong/1/1968. rHK-17R contained the single point-mutation H17R in the HA, which decreased its conformational stability and increased pH threshold of fusion. Thus, 50 % of rHK-wt HA underwent irreversible conformational transition at pH 4.8, whereas in the case of rHK-17R the 50 %-transition was achieved at pH 5.3 (Gerlach et al. 2017). After expansion, the identity of the HA was confirmed by sequencing, the viruses were purified, aliquoted and stored at -80 °C. Viral titres were determined using single-cycle infection assay as described previously (Matrosovich et al. 2007). In brief, MDCK cells were inoculated with serially diluted virus and incubated for 18 h in the absence of trypsin. The cells were fixed, permeabilized, and stained for nucleoprotein expression with primary mouse anti-influenza A virus nucleoprotein antibody (9G8, 0.5 mg/ml, Abcam). For visualization of NP-positive cells, a secondary peroxidase-labelled anti-mouse antibody was added, followed by washing and detection of peroxidase activity with KPL True Blue™ substrate (SeraCare). Stained infected cells were counted under the microscope.

### 2.3 Infection media

Media used for infections (below, “infection media”), such as DMEM (Pan BioTech, P04-036001) and RPMI (Pan BioTech, P04-18000), were always supplemented with 0.1 % BSA (Gibco), 2 mM L-glutamine (Invitrogen) and 100 IU/ml penicillin-streptomycin (Invitrogen). Commercial OptiMEM medium (Gibco, 31985070) already contained L-glutamine and was only supplemented with 0.1 % BSA and 100 IU/ml penicillin-streptomycin. Infection media did not contain trypsin in order to limit influenza infection to one cycle.

As controls, we used samples of infection medium that were not supplemented with L-glutamine, but contained all other mentioned supplements. Additionally, we used samples of infection medium that were supplemented with 1× GlutaMAX™ (Gibco, 35050061) instead of L-glutamine. To control for the effect of HEPES, we used DMEM infection medium with above-mentioned supplements and added 5958 mg/L HEPES (Sigma). We also used OptiMEM infection medium and added 2979 mg/L HEPES, as commercial OptiMEM already contains HEPES at half of the concentration in RPMI medium (2400 mg/L) (Thermo Fisher Scientific, DE, 2020).

Aliquots of infection media were stored at either i) 4 °C for four days, ii) 37 °C for four days, or iii) 4 °C for at least three months. The first condition represented fresh and adequately stored infection medium. The second was expected to accelerate temperature-induced decomposition of L-glutamine. The third was used to represent “old” infection medium.

### 2.4 Infections

0.125 × 10^6^ MDCK cells per well were seeded on 48-well plates in growth medium and were allowed to adhere overnight. Cells from one well were harvested and counted. The other wells were washed two times with calcium- and magnesium-containing PBS (DPBS^++^, Pan BioTech, P04-35500) and then supplied with infection medium. All infection media had been brought to room temperature prior to application. The cells were inoculated with purified virus at 3 MOI and were incubated for eight hours at 37 °C, 5 % CO_2_.

Eight hours post infection (hpi) cells were washed twice with PBS containing no calcium and magnesium (DPBS_def_, Pan BioTech) and detached with trypsin (Gibco). Enzymatic detachment was stopped by resuspending the cells in growth medium. The suspension was transferred to FACS tubes cells were washed twice with FACS-buffer (DPBS_def_ + 3 % foetal calf serum (Sigma-Aldrich) + 2 mM EDTA (Roth) + 0,001 % sodium azide (Roth)). Cells were fixed with paraformaldehyde at a final concentration of 2 % for 20 minutes at room temperature. In order to make intracellular viral protein accessible, the cells were permeabilized for 30 minutes with saponin-buffer (FACS-buffer + 0.5 % saponin from *quillaja bark* with sapogenin content ≥10 % (Sigma)). Cells were washed twice and incubated with primary mouse anti-influenza A virus nucleoprotein antibody (9G8, Abcam, 0.5 mg/ml) at a final concentration of 1:500 for 1 hour at 4 °C, followed by two washings with FACS-buffer. Alexa Fluor 633 goat anti-mouse IgG (H+L) cross-adsorbed antibody (Invitrogen, 2 mg/ml) was used as a secondary antibody at a final concentration of 1:500. Incubation with this antibody was performed for 1 hour at 4 °C. The cells were washed twice, resuspended in 100 μL FACS-buffer and analysed using FACS Calibur® and FlowJo (Becton Dickinson). Cells displaying morphology of living cells in forward/sideward scatter (FSC-SSC) analysis were selected and gated for an allophycocyanin-positive (APC^+^) signal. The percentage of APC^+^ cells served as a readout for infectivity.

### 2.6 Infection inhibition by ammonium chloride

Infections of MDCK cells with rHK-wt and rHK-17R were studied as described above using DMEM or RPMI infection medium containing serial two-fold dilutions of ammonium chloride with concentrations from 2 to 0.125 mM of NH_4_Cl. Infectivity was analysed as described above.

### 2.7 Ammonium quantification

Total ammonium was quantified colorimetrically using Ammonium Assay Kit (Abcam, 83360) following the manufacturer’s protocol. In brief, we prepared DMEM infection medium with the supplements as explained above (see 2.3), using DMEM deficient in both phenol red and pyruvate (Pan BioTech P04-01161), as these substances interfere with the assay. Storage was conducted at either 4 °C for four days (fresh medium), 37 °C for four days (pre-warmed medium) or 10 days at ambient temperature (old medium). Media were pre-diluted 1:10 in the assay buffer and mixed with an equal volume of the assay’s enzyme master mix. Absorbance at 570 nm was measured using a microplate reader.

### 2.9 Statistics

Data were analysed using GraphPad Prism 10. The statistical methods are mentioned in the figure legends. Differences characterized by *p*-values greater than 0.05 were considered as not significant.

## 2. Results

### Inadequately stored DMEM inhibits infectivity of rHK-wt

We firstly addressed the question whether the choice of cell culture infection medium and its storage can affect IAV infectivity. The three widely used media DMEM, RPMI, and OptiMEM, each containing L-glutamine, were subjected to three different storage conditions. Storage at 4 °C for four days represented adequately stored medium (below, “fresh medium”), expected to accumulate minimal concentrations of L-glutamine-derived ammonium. Storage at 37 °C for four days represented laboratory handling, where researchers pre-warm their media in a water bath prior to their experiments or keep media at 37 °C to test for contaminations (below, “pre-warmed medium”). Storage at 4 °C for at least three months represented inadequately long storage (below, “old medium”). MDCK cells were infected with the two recombinant IAVs rHK-wt or rHK-17R at 3 MOI. The viruses differed solely in the pH optimum of fusion of their HA protein, i.e. rHK-wt having its optimum at pH 4.8 and rHK-17R having its optimum at pH 5.4 (Gerlach et al. 2017). Viral infectivity was quantified via immunostaining for viral NP expression in MDCK cells. We observed pronounced differences in the infection efficacy of the viruses, which strongly depended on both the culture medium and its storage conditions (Fig. 1). Fresh DMEM with L-glutamine was found to allow for infections of near-to-identical efficacy (80 %) for both viruses (Fig. 1A, left panel). Interestingly, infectivity of rHK-wt was 4-fold reduced in old DMEM when compared to its pH-labile counterpart rHK-17R. A less prominent but still significant difference was detected in pre-warmed medium. In this case, the pH-stable virus rHK-wt infected over 2-fold less efficiently than rHK-17R.

**Figure 1:**
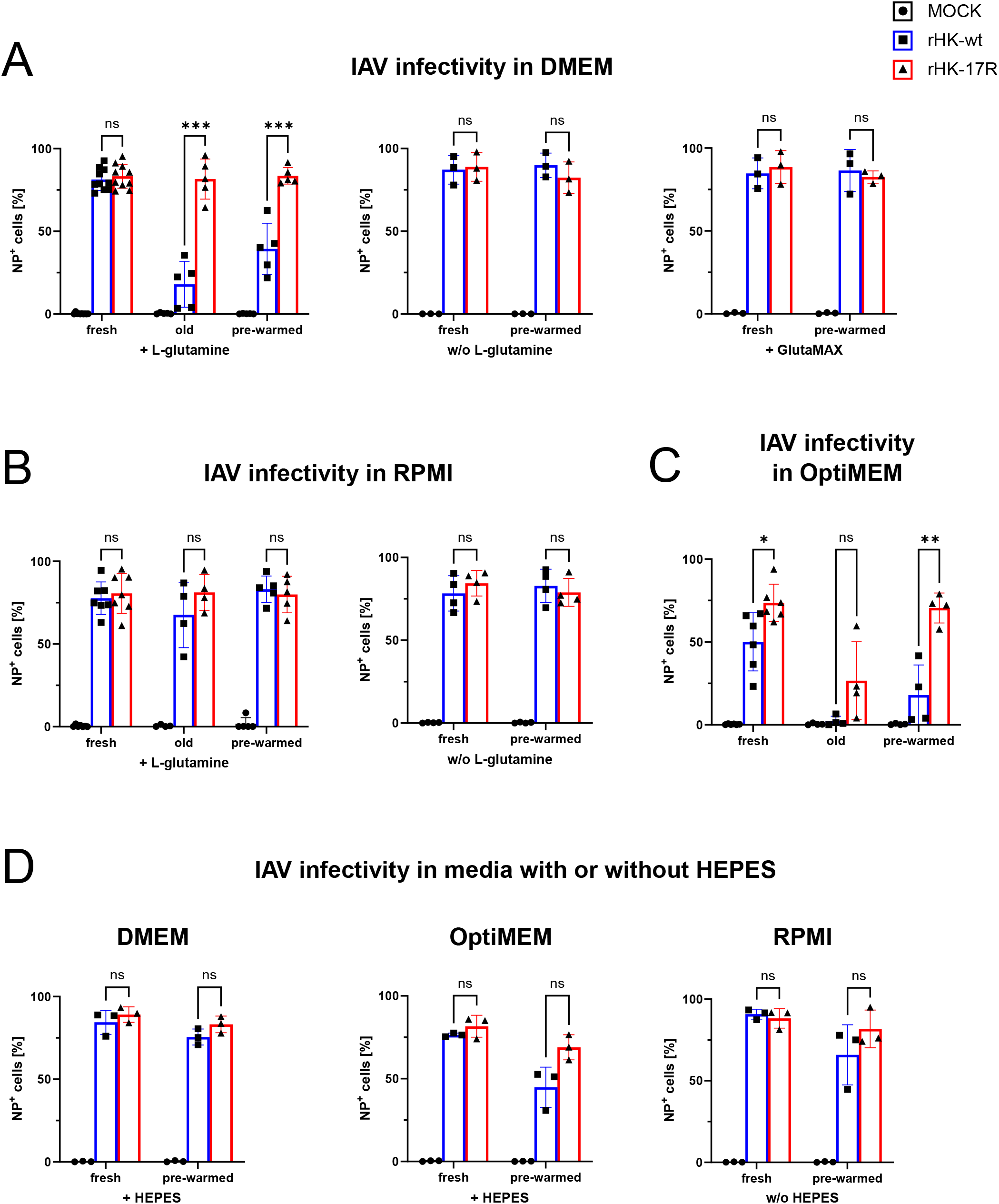
IAV infectivity depends on culture medium and correlates with the presence of L-glutamine. Infectivity of rHK-wt and -17R was quantified in infections of MDCK cells at 3 MOI, performed in one of three cell culture media with various supplements as indicated. Media were stored for four days at 4 °C (fresh), four days at 37 °C (pre-warmed) or for at least three months at 4 °C (old). (A) Quantification of infectivity in DMEM with L-glutamine (left panel), without L-glutamine (middle panel) or with GlutaMAX (right panel). (B) Quantification of infectivity in RPMI with L-glutamine (left panel) or without L-glutamine (right panel). (C)Quantification of infectivity in OptiMEM, which received L-glutamine during its production. (D)Quantification of infectivity in media with or without HEPES. Single data points indicate percentages of NP^+^ cells from independent experiments (n ≥ 3), measured via flow cytometry. Bars represent means ± standard deviations. Differences in infectivity of the viruses in cell culture medium were compared by unpaired t tests using Holm-Šídák correction for multiple comparisons. Asterisks indicate calculated p-values (ns, p > 0.05; *, p < 0.05; **, p < 0.01; ***, p < 0.001).

L-glutamine is known to decompose in aqueous solution, producing ammonium ions (Tritsch and Moore 1962). To test whether L-glutamine-derived ammonium was responsible for the observed differences in IAV infectivity, we used medium deficient in L-glutamine. Indeed, both viruses infected MDCK cells efficiently and independent of the storage conditions (fresh or pre-warmed) when no L-glutamine was added to the medium (Fig. 1A, middle panel). Considering that the viruses were equal in all properties except for their pH optimum of membrane fusion, these observations imply that decomposition of L-glutamine, generation of ammonium and its effect on endosomal acidification were responsible for the inhibition of cell entry by rHK-wt. In contrast, a lack of significant inhibition of rHK-17R agreed with the ability of its HA to mediate fusion at a relatively high endosomal pH.

In some commercial media, L-glutamine is substituted by L-glutamine-containing dipeptides, which are more stable in aqueous environments. Arii and co-workers calculated that the shelf life of L-alanyl-L-glutamine is 5.3 years at 25 °C, and 7.1 months at 40 °C (Arii et al. 1999). Li and colleagues found that the ammonium-induced effects in their experiments did no longer occur when they substituted L-glutamine by L-alanyl-L-glutamine (GlutaMAX) (Li et al. 2016). In this view, we prepared DMEM containing GlutaMAX instead of L-glutamine. The GlutaMAX-containing medium was stored under the conditions described above. rHK-wt and rHK-17R infected MDCK cells with comparable efficiency in fresh DMEM with GlutaMAX (Fig. 1A, right panel), and infection levels did not differ from levels observed in fresh L-glutamine-containing DMEM (Fig. 1A, left panel). Importantly, we were not able to detect any differences between infection efficiencies of these two viruses in pre-warmed medium (Fig. 1A, right panel). These data put further emphasis on the hypothesis that the decay of L-glutamine is responsible for the HA-stability-dependent viral attenuation in DMEM.

### Improper storage of RPMI has no significant effect on infectivity of IAVs

RPMI medium is commonly used for cultivation of peripheral blood mononuclear cells (PBMCs), and we recently studied infection of PBMCs by IAVs in this medium (Dorna et al. 2022). In this view, we investigated infections with both IAVs conducted in RPMI, which was stored under the conditions explained above. In contrast to DMEM, storage and pre-warming of the original RPMI infection medium and RPMI deficient in L-glutamine did not affect infectivity of rHK-wt or rHK-17R (Fig. 1B). These data suggest that RPMI preserves reproducibility of infections and does not exert differential effects on infectivity of IAVs differing by HA-stability as they were observed in DMEM (Fig. 1A).

### IAV infectivity is decreased in fresh OptiMEM compared to fresh DMEM and RPMI

Cell culture media that receive L-glutamine during production may accumulate ammonium due to sub-optimal storage or shipping conditions before their arrival in the laboratory (Li et al. 2016). We assayed whether infections were impaired in OptiMEM, a medium which is supplemented with L-glutamine during its manufacturing and is shipped at ambient temperature (Thermo Fisher Scientific, DE, 2020). To our surprise, infectivity in fresh OptiMEM was impaired (Fig. 1C) relative to the infectivity observed in DMEM and RPMI (Figs. 1A, B). We detected a significant difference in infection efficiency of rHK-wt versus rHK-17. The difference was further increased by using pre-warmed OptiMEM, where rHK-17R infected almost 4-fold more cells than rHK-wt, while rHK-17R remained unaffected (Fig. 1C). Notably, long-term storage completely inhibited rHK-wt and reduced the infectivity of rHK-17R by almost 3-fold. Together, the attenuation of the pH-stable virus rHK-wt in fresh OptiMEM demonstrates a pH-stability-directed effect that can be enhanced by excessively long storage and by storage at elevated temperature.

### HEPES counteracts inhibition of virus infectivity in pre-warmed and old media

The release of ammonium from L-glutamine is known to depend, in part, on ionic strength and pH of the medium (Tritsch and Moore 1962). Assuming that supplements present in DMEM and RPMI could affect the degradation kinetics of L-glutamine, we focused on the differences in the components used as buffers. HEPES is included as a buffer in RPMI, while DMEM is buffered by sodium bicarbonate. To test if HEPES could be responsible for the observed stability of stored RPMI (Fig. 1B), we used normal L-glutamine-containing DMEM infection medium and supplemented it with 5958 mg/L HEPES in accord with the concentration of HEPES in RPMI. Fresh or pre-warmed HEPES-supplemented DMEM was then used for infections with rHK-wt and rHK-17R. HEPES had no influence on the infectivity of either virus in the fresh medium (Fig. 1D, left panel). Strikingly, no reduction of infectivity of rHK-wt was observed in pre-warmed DMEM with HEPES, in contrast with the loss of infectivity in the pre-warmed DMEM without HEPES (Fig. 1A).

We next investigated if infections conducted in OptiMEM can be rescued by increasing the concentration of HEPES in OptiMEM, in which the HEPES concentration is half of that in RPMI (2400 mg/L) (Thermo Fisher Scientific, DE, 2020). We observed that the difference in infectivity between the viruses in fresh OptiMEM was no longer detectable (Fig. 1D, middle panel). In pre-warmed OptiMEM, we continued to observe a disparity between the percentages of infected cells, but the difference was not as significant as in the original OptiMEM (Fig. 1C). As a control, we conducted infections in RPMI medium containing no HEPES. As expected, the use of fresh RPMI without HEPES allowed for similar infection levels of both viruses (Fig. 1D, right panel). Surprisingly, pre-warming did not lead to a statistically significant difference between the infectivity of the two viruses. In summary, in our experiments, a higher concentration of HEPES facilitated IAV infections and reduced the advert effect of medium storage on IAV infectivity. However, as the removal of HEPES from RPMI did not restore a disparity between the two viruses following pre-warming of media, our results imply that HEPES is not the sole substance responsible for the observed stability of RPMI-based infection medium during storage.

### Virus-neutralizing effect of ammonium ions depends on stability of viral HA

In order to assay how ammonium ions affect the infectivity of the two IAVs used in this study, we added ammonium chloride (NH_4_Cl) in two-fold serial dilutions ranging from 2 to 0.125 mM to DMEM and RPMI and used these media for infections of MDCK cells with either the pH-stable virus rHK-wt or the pH-labile virus rHK-17R. We observed that rHK-wt was more susceptible to inhibition by NH_4_Cl than rHK-17R (Fig. 2). In both tested media, 1 mM NH_4_Cl completely inhibited infections by rHK-wt, while more than 50 % cells were infected with rHK-17R at the same concentration of NH_4_Cl. rHK-wt restored its normal infectivity at approximately 0.125 mM NH_4_Cl, and both viruses infected with comparably efficiency in the absence of NH_4_Cl. These experiments demonstrated that under conditions of our infectivity assay ammonium ion concentrations below 2 mM significantly reduced the infectivity of pH-stable IAVs with a notably lower inhibitory effect on the pH-labile variant.

**Figure 2:**
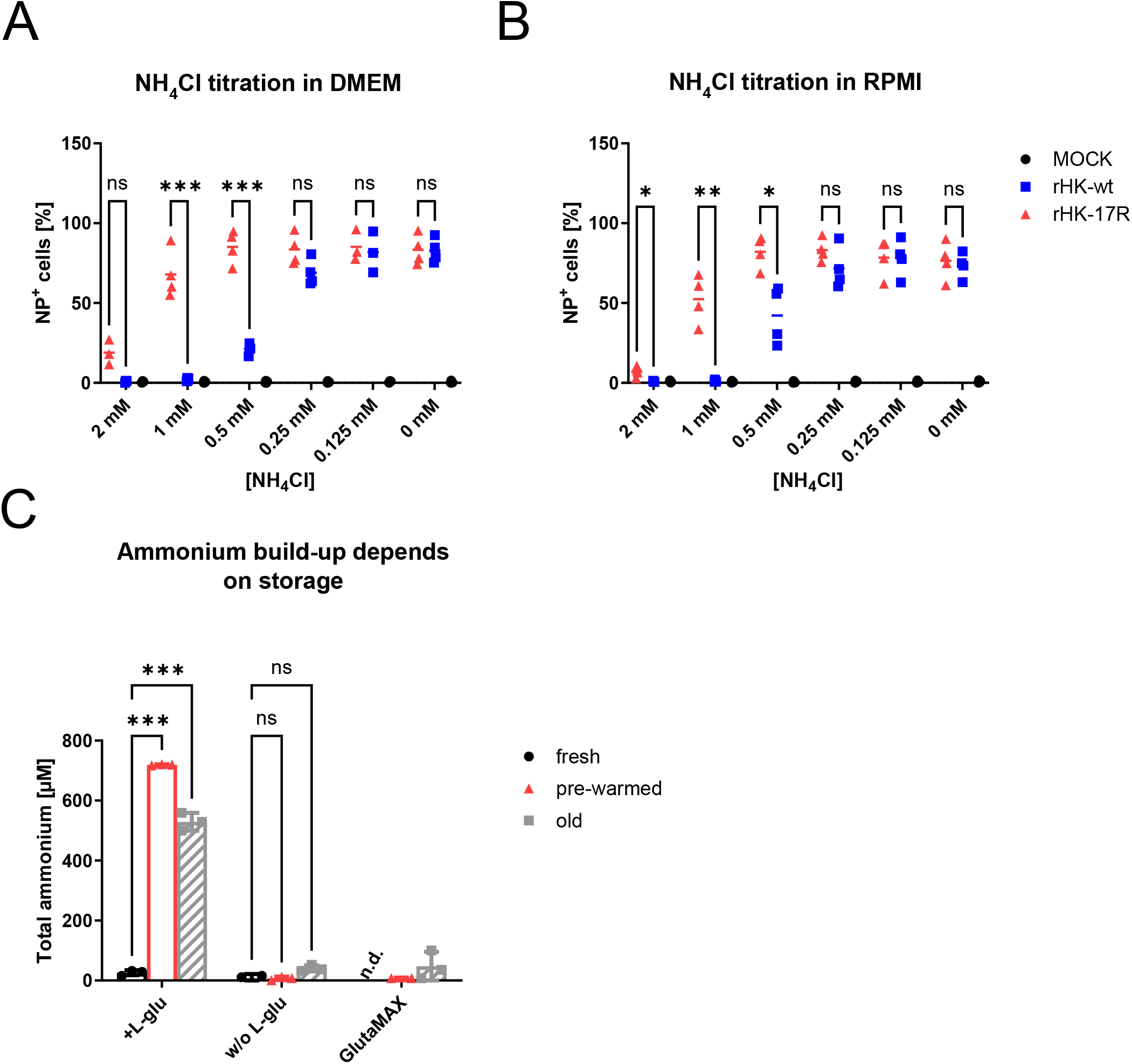
Titration of IAV infectivity against exogenously added NH_4_Cl and quantification of ammonium build-up. (A) MDCK cells were infected with rHK-wt or rHK-17R at 3 MOI in DMEM-based medium containing 0.125 mM to 2 mM NH_4_Cl. Infections in medium without NH_4_Cl and uninfected cells served as a control. Infections were analysed as described in Figure 1 legend. (B) The assay as in (A) but using RPMI instead of DMEM. Single data points indicate percentages of NP^+^ cells from independent experiments (n = 4). Medians are shown. Percentages of the cells infected by individual viruses were compared by unpaired t tests using Holm-Šídák correction for multiple comparisons. Asterisks indicate calculated p-values (ns, p > 0.05; *, p < 0.05; **, p < 0.01; ***, p < 0.001). (C) Quantification of total ammonium accumulation using a commercial kit. DMEM infection medium was prepared with or without L-glutamine (L-glu) or with GlutaMAX. Medium batches were stored at 4 °C for four days (fresh), at 37 °C for four days (pre-warmed) or at room temperature for 10 days (old). Single data points indicate results from three independently prepared medium batches. Measurements below the detection limit are indicated by “n.d.” (not detected). Bars show means ± standard deviations. Data were compared by two-way ANOVA with Tukey correction. Asterisks in figure (A) and (B) indicate p-values for differences in counts of positive cells of rHK-wt vs. rHK-17R. Asterisks in figure (C) indicate differences in results between medium groups as indicated (ns, p > 0.05; *, p < 0.05; **, p < 0.01; ***, p < 0.001).

### Release of ammonium in improperly stored DMEM medium

In order to validate our findings, we quantified the release of ammonium in DMEM with or without L-glutamine or with GlutaMAX. Medium batches were subjected to 4 °C for four days (fresh medium), 37 °C for four days (pre-warmed medium), or ten days at ambient temperature (old medium). Of note, the latter conditions have been described by Li and colleagues to result in a complete degradation of L-glutamine (Li et al. 2016). We detected over 20-fold more ammonium in pre-warmed and old medium compared to fresh medium (Fig. 2C). Remarkably, media without L-glutamine or with GlutaMAX contained minimal amounts of ammonia that were close to the detection limit, irrespective of the storage conditions. We here observed that prolonged medium storage or medium pre-warming released significant concentrations of ammonium. Our data demonstrate that L-glutamine was a source of ammonium in concentrations sufficient to inhibit infectivity of pH-stable IAVs.

## 3. Discussion

In this study, we demonstrated that ammonium emerges rapidly in L-glutamine-containing cell culture media under conditions that are typical for laboratory practice. We found that the use of inadequately stored media can inhibit single-cycle infections of influenza viruses in cell culture, which in its turn will have implications for virus multicycle replication and production. Since we found that the effect depended on properties of the viral HA, and as the release of ammonium is poorly controlled, using improperly stored media can impair reproducibility of results.

Detrimental effects of ammonium on various in vitro settings were comprehensively reviewed by Schneider (Schneider 1996). With regard to influenza viruses, early work revealed ammonium-related reduction of viral replication (Eaton and Scala 1961; Jensen et al. 1961), which was later explained by ammonium-mediated inhibition of viral pH-dependent fusion in the endosomal compartment (Matlin et al. 1981; Doms et al. 1986). However, some reports used up to 10 mM and higher concentrations of exogenously added ammonium (Whittaker et al. 1996; Morris et al. 1999), which is notably more than the maximum ammonium concentration that can be generated from L-glutamine decomposition in mammalian cell culture media, which corresponds to the typically used 2-4 mM of L-glutamine. Reports on the inhibition of influenza viruses by ammonium released from L-glutamine are ambiguous. For example, experiments performed by Genzel and colleagues did not detect inhibition of influenza virus production by ammonium emerging from L-glutamine metabolism or decomposition (Genzel et al. 2004). Most studies agree in the conclusion that generation and/or adverse effects of ammonium depend on both the type of cells and virus strains used (Schneider 1996; Jagušić et al. 2016; Decrey et al. 2016). Our results highlight the role of HA conformational stability in susceptibility of IAVs to ammonium-mediated inhibition in cell culture, and they may explain apparent discrepancies between reported effects of ammonium on influenza viruses.

We observed that effects of cell culture media storage on IAV infection depended on the type of the medium. While infections in improperly stored DMEM and OptiMEM were prone to significant inconsistencies, storage had little if any effect on IAV infection in RPMI (Fig. 1). As all tested media were subjected to identical storage conditions, we conclude that they differed by the rate of ammonium accumulation during storage and this effect depended on composition and pH of the medium (Tritsch and Moore 1962). We found that addition of HEPES to DMEM and OptiMEM reduced deterioration of these media during storage (Fig. 1D). We hypothesize that several mechanisms can be responsible for this effect of HEPES. On one hand, HEPES may reduce decomposition of L-glutamine and thus production of ammonia by affecting the pH of the medium (Tritsch and Moore 1962). On the other hand, HEPES-mediated effects on pH may alter the protonation state of liberated ammonia. In the aqueous, mildly alkaline environment of cell culture media, ammonium exists mainly in its protonated form NH_4_^+^ (Schneider 1996), which cannot diffuse across membranes, and there is only a low level of uncharged, lipophilic NH_3_, which can enter endosomes through undirected diffusion. Thus, increasing the portion of positively charged molecules by changes in medium pH could reduce the share of molecules that can enter endosomes, and thereby affect endosomal acidification.

HEPES is taken up by cells (Depping and Seeger 2019) and evidence was gathered that HEPES is not innocuous (Bowman et al. 1985; Douglas et al. 1993; Cook et al. 2020; Tol et al. 2018). In particular, Tol and colleagues showed that HEPES activates transcriptional events which promote endosomal biogenesis (Tol et al. 2018). Therefore, in addition to its buffering activity, other infection-facilitating effects of HEPES must be considered.

Our results demonstrate that decomposition of L-glutamine and production of ammonia in cell culture media may reduce infectivity and hamper propagation of IAVs. Viruses with a pH-stable HA, such as human viruses and viruses of wild aquatic birds, will be particularly sensitive to inhibition. It is therefore advisable to either use L-glutamine-deficient media or substitute L-glutamine by stable analogues, such as GlutaMAX. Prolonged and inadequate storage of L-glutamine-containing medium should be avoided.

There is a growing interest in studying membrane fusion properties of influenza viruses and their role in replication and tropism (Russell et al. 2018). Inhibition of IAV infection in cell culture by ammonium salts represents one of the assays used to compare conformational stability of the HA and fusion pH optimum of viral isolates (Doms et al. 1986; Koerner et al. 2012; Gerlach et al. 2017). Our findings should alert researchers about potential presence of lysosomotropic agents in cell culture media, which may affect assay results and thus compromise adequate comparison of different IAVs.

## Acknowledgements

This work was supported by Deutsche Forschungsgemeinschaft (DFG, German Research Foundation) Project-ID 369799452 – TRR237 - A02 to SB and Project-ID 197785619 - SFB 1021 to MM. We want to thank Thomas Gerlach and Luca Hensen for the production of the recombinant influenza viruses used in this study.

## Author contributions

NBK, AK performed the experiments. JD, SB, NBK, AK, MM designed the analyses. JD, SB, AK conceived and supervised the study. JD contributed analysis tools. NBK, SB wrote the initial manuscript. MM, JD, AK reviewed the manuscript.

